# Nucleosomal embedding reshapes the dynamics of abasic sites

**DOI:** 10.1101/2020.02.26.966366

**Authors:** Emmanuelle Bignon, Victor Claerbout, Tao Jiang, Christophe Morell, Natacha Gillet, Elise Dumont

## Abstract

Apurinic/apyrimidinic (AP) sites are the most common DNA lesions, which benefit from a most efficient repair by the base excision pathway. The impact of losing a nucleobase on the conformation and dynamics of B-DNA is well characterized. Yet AP sites seem to present an entirely different chemistry in nucleosomal DNA, with lifetimes reduced up to 100-fold, and the much increased formation of covalent DNA-protein cross-links, refractory to repair. We report microsecond range, all-atom molecular dynamics simulations that capture the conformational dynamics of AP sites and their tetrahydrofuran analogs at two symmetrical positions within a nucleosome core particle, starting from a recent crystal structure. Different behaviours between the deoxyribo-based and tetrahydrofuran-type abasic sites are evidenced. The two solvent-exposed lesion sites present contrasted extrahelicities, revealing the crucial role of the position of a defect around the histone core. Our all-atom simulations also identify and quantify the occurrence of several spontaneous, non-covalent interactions between AP and positively-charged residues from the histones H2A and H2B tails that prefigure DNA-protein cross-links. This study paves the way towards an in silico mapping of DNA-protein cross-links.

## INTRODUCTION

Apurinic/apyrimidinic (AP) sites are the most frequent spontaneous DNA lesions, amounting to 5,000–10,000 lesions per day per mammal cell (1, 2). They are generated by hydrolysis of the N-glycosidic bond between a nucleobase and deoxyribose (3), spontaneously or as intermediates in the excision repair of modified bases. Their repair by dedicated enzymes turns out to be most efficient by base excision repair (BER) but also nucleotide excision repair (NER) (4) yet accumulation of AP sites, notably triggered by exogenous oxidative stress, represents a threat for genome integrity. Notably AP sites can act as precursors towards much more deleterious lesions such as strand breaks and DNA-proteins cross-links (DPC, see Figure 1-A rightside) (5), through the reactive aldehyde moeity of the open conformation (6) (see Figure 1-a). AP sites chemistry is hence of utmost importance for the development of anticancer therapies (7). Structural and thermodynamical (8) features of AP sites as well as their repair (9) have been intensively studied for oligonucleotides systems and G-quadruplexes (10): the structural reorganization of the “hole” left within the B-helix, the non-covalent interactions developed by the orphan nucleobase on the opposite strand remains to be elucidated. Recent chemical methods have been proposed to sequence them in DNA with single-nucleotide resolution (11). Yet until recently, insights into the chemistry of AP sites within nucleosomal DNA were scarce. Greenberg and coworkers have reported vastly shortened lifetimes for AP sites embedded within nucleosome core particles (NCP). A recent structure, shown in Figure 1-B, has been obtained by Osakabe and coworkers (12). This structure features a segment of ds-DNA, comprising 145 base pairs (bp), wrapped onto an octameric core of two copies of four types of histone proteins, H2A, H2B, H3 and H4. It harbors two tetrahydrofuran (THF) analogs of AP sites at symmetrical superhelical position SHL4.5.

**Figure 1.**
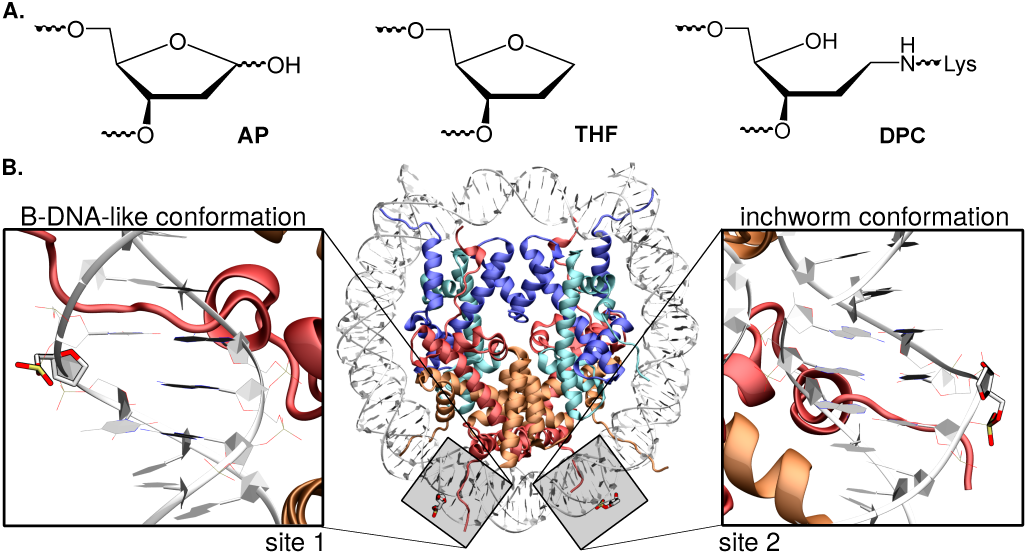
A. Chemical structure of an AP site, its THF analog, and DNA-protein crosslink (DPC) formed upon reaction between lysines and AP sites. B. Cartoon representation of the crystal structure of a NCP harboring two symmetrical, solvent-exposed tetrahydrofuran residues (sites 1 and 2; PDB ID 5JRG). Site 1 exhibits a classical B-DNA-like conformation, whereas site 2 adopts the so-called inchworm conformation following the exclusion of THF and an adenine on the opposite strand.

The histones possess short tails which play a significant role, notably their interactions with AP sites give rise to a much enhanced reactivity compared to naked DNA (100-fold more reactive), with a strong dependence on the exact positioning of the AP site on the NCP (13, 14).

Molecular dynamics has become a dominant method to shed complementary insights on: structural (15) and mechanical properties (16), reactivity and repair of AP sites (17, 18, 19), and effects of clustered AP sites (20, 21). Also a few all-atom molecular dynamics simulations have been performed on NCPs, undamaged (22, 23, 24), stacked (25), with double-strand breaks (26) or to decipher the impact of acetylation of histone tails (27). As it has become possible to reach the microsecond range for such systems (28), they can serve as a computational microscope (29) to probe the conformational and mechanical cross-talk between an abasic site and its N-DNA environment. In this study, microsecond range all-atom simulations were performed to unravel the structural outcome and time evolution of AP-containing and THF-containing NCP, starting from the X-ray structure obtained by Osakabe *et al.* (12).

## MATERIALS AND METHODS

The crystal structure of a NCP harboring symmetrical THF experimentally invoked as AP sites analogs was used as starting structure for our models (12). Two starting models were generated from the crystal structure: one with THF as initially in the crystal, the other one with an abasic site. The missing adenine on the DNA strand facing the damaged site exhibiting the ‘inchworm’ conformation shown in Figure 1-B was added *in silico* using `xleap`. Crystallographic water molecules and chloride ions were kept, and potassium ions were added to neutralize the system. Protonation states of histone core residues were assigned using propka3.1 (30), and histidines were protonated at the *ε* position. The two 15-bp sections harboring each damaged sites (1 and 2) were extracted from the crystal structure and used to run reference simulations on control B-DNA structures with either Ap or THF - see Fig. 1. Each system was soaked in a TIP3P water box (truncated octahedron) with a buffer of 10 Å, resulting in ~73,500 atoms for NCPs and ~25,000-35,000 atoms for the reference 15-bp oligonucleotides. The ff14SB force field was used in our simulations (31), together with the OL15 corrections for nucleic acids (32, 33). Force field parameters for non-canonical lesions were generated for THF and taken from previous works for AP sites (18, 21) - see Table S1 and Figure S1. All MD simulations were performed with the Amber and Ambertools 2018 packages (34). The NCP starting structures were carefully optimized through four 10,000 steps energy minimization runs with restraints on the histones and DNA residues, gradually decreased from 20 to 0 kcal.mol^*−*1^.Å^2^. The system was then heated from 0 to 300K in a 60ps thermalization run. The temperature was subsequently maintained at 300K along the simulation using the Langevin thermostat with a collision frequency *γ*ln of 1 ps^*−*1^. The system was then equilibrated during 2ns in the NPT ensemble. Finally, a one microsecond production run was carried out. For the two NCP starting structures (AP and THF), three simulation replicates (MD1, MD2, MD3) were generated using different starting velocities. The Particle Mesh Ewald method with a 9.0 Å cutoff was used to account for electrostatic interactions. Production runs for the B-DNA reference structures were 1 *µ*s long. Post-processing and structural analyses were carried out using AmberTools 2018 and Curves+ (35) programs. Analyses were performed on the 13-bp section surrounding the damaged sites as showed in Figure 1. Analyses of the 15-bp DNA reference simulations were consistently ran on the 13-bp to avoid any edge effects from the floppy first and last base pairs of the oligonucleotide. Clustering of MD ensembles was carried out according to the RMSD of the 13-bp harboring the damaged site showed in Figure 1-B.

The sheer size of the dimensionality of the data from molecular dynamics simulations often makes it difficult for visual inspection of the trajectory, especially for such all-atom N-DNA trajectories. Principal components analysis (PCA) can help reducing the dimensionality by reconstructing a new configurational space that contains the most important degrees of freedom, providing a better angle for grasping the essence of the chemical process. In practice, the PCA method creates a covariance matrix from the coordinates of the trajectory as input. A set of eigenvectors are then obtained by diagonalizing the covariance matrix, serving as basis of a new configurational space, with each of them being a direction of motion. The magnitudes of the eigenvalues indicates the data variance in each of these directions of motions, the eigenvector with the largest eigenvalue is called the first principal component, and so on. The PCA was performed with a home brew script utilizing the Scikit-learn (36) library, the internal coordinates (inverse distance between geometric centers of two residues) of the trajectory as input instead of Cartesian coordinates, due to better performance (37). To obtain the per residue importance, the sum of the weighted principal components up to certain threshold with the corresponding as weights is taken. The sum was then normalized and mapped back to the residues.

## RESULTS

The nucleobases numbering used corresponds to the labelling shown in Figure 2-B.

**Figure 2.**
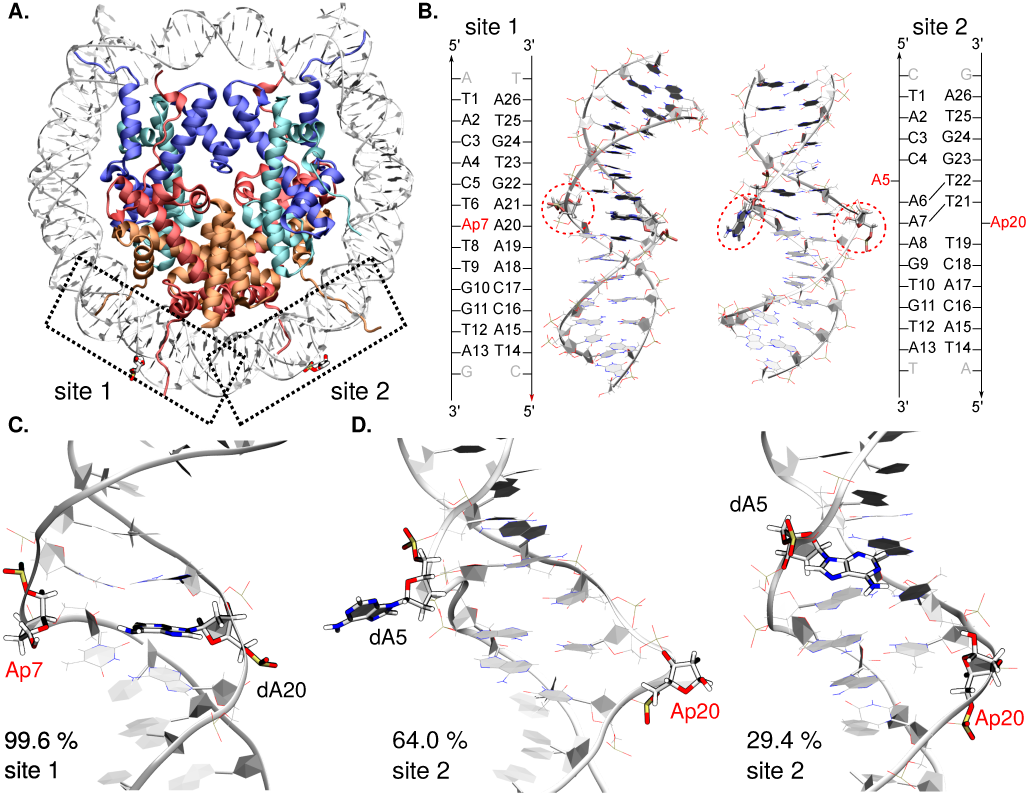
**A.** Structure of the THF-containing NCP structure (PDB ID 5JRG (12)). Histones H2A, H2B, H3, and H4 appear in red, orange, blue, and cyan, respectively. The two symmetric lesion sites are depicted in licorice, site 1 on the left and site 2 exhibiting the inchworm conformation on the right. **B.**Cartoon representation of the 15-bp sections harboring the lesion sites, and their corresponding sequences. The damaged sites (here AP), as well as the *in silico* modeled dA5 for site 2, are encircled in red. Structural analyses were performed on these 15-bp sections only. Reference MD simulations of naked DNA were performed on the same sequences, with one extra base pair at each extremity to avoid strong end effects on the structural parameters. **C.** and **D.** Representative structures of the major clusters and percentage of occurrence for site 1 (C.) and site 2 (D.) AP-containing NCP. The lesions sites are depicted in licorice, as well as the facing base for site 1 and the ejected dA5 for site 2. Conformations sampled for site 1 mostly belong to one dominant cluster (99.6%) while site 2 exhibits two main different conformations.

### The nucleosomal embedding impacts the bend angles uniformly along the histone core

Molecular modeling techniques were employed to probe the structural behavior of NCP harboring deoxyribose- and tetrahydrofuran-type abasic sites, hereafter referred as AP and THF. Three unbiased classical MD simulation replicates of 1*µ*s each (MD1, MD2, and MD3) were carried out on the NCP structure exhibiting either AP sites or THF analogs at symmetrical superhelical location (SHL) of 4.5, as positioned in the crystal structure by Osakabe *et al.* (12). We describe hereafter the local structural features of the two 15-bp sections embedding the damaged sites 1 and 2, as shown in Figure 2. A first overall measure of the structural impact of an abasic site (or more generally a DNA lesion) on the B-helicity is the bend angle, which was monitored along our MD simulations. For AP sites in the NCP, the bend angle remains stable around 55 ± 9° vs. 27 ± 13° for the control AP-containing sequence (see Figure S1). The bend angle reflects the mechanical constraint exerted along the histone core as the DNA sequence wraps around it (38), and is found to be rather constant including in the vicinity of AP sites. However the higher curvature may impact on the ease to flip out the AP site or a proximal nucleobase, which lead us to investigate more locally the structural distortions in the vicinity of the AP sites.

### Extrahelicity differs at sites 1 and 2

Our simulations allow to delineate significant differences of local deformation between AP sites. Extrahelicity of AP sites is a most important structural signature that can also be monitored along our simulations (18, 21). The extrahelicity criteria is based on the distance between the AP site’s C1’ atom and C1’ of its facing nucleotide, the threshold value above which the AP site is considered as extrahelical being 14Å (39). The intrinsic propensity for an AP site to flip out the B-helix within an oligonucleotide is typically of 30% (16), as is indeed observed in the simulations of the control 15-bp sequences. Averaged extrahelicity values reported in Table 1 highlight two very different behaviors for AP sites at site 1 and site 2 within the NCP, with respective values of 19.0% and 88.2%. The NCP environment can induce either a slightly lower extrahelical character at site 1 and, on the contrary, an exacerbated flipping of the lesion at site 2. The initial inchworm conformation of site 2, as well as the ejected dA5 on the facing strand might favor the extrahelical character of the lesion site, which is maintained along our simulations. Noteworthy, the same trend is observed for THF-containing NCP (average average 5.3% site 1 and 85.3% site 2), yet the much higher extrahelicity exhibited by THF in the control 15-bp (42.7 % and 72.8%).

**Table 1.** Detailed and average of the percentage of extrahelicity in the three 1 *µ*s MD replicates for AP and THF analogs at site 1 and site 2 in the NCP, and in the 1 *µ*s 15-bp control simulations.

| System | all | MD1 | MD2 | MD3 | Control 15-bp |
| --- | --- | --- | --- | --- | --- |
| AP (Site 1) | 19.0% | 14.4 % | 12.8 % | 29.7 % | 32.1 % |
| AP (Site 2) | 88.2% | 67.0 % | 99.6 % | 97.9 % | 39.1 % |
| THF (Site 1) | 5.3 % | 3.5 % | 1.0 % | 11.5 % | 42.7 % |
| THF (Site 2) | 85.3% | 56.6 % | 99.5 % | 99.7 % | 72.8 % |

### Cluster analysis of the MD ensembles

Our thorough structural analysis along the MD ensembles reveals a complex structural behavior with several regimes for the AP-sites conformation as shown for main representative clusters of the three MD replicates at sites 1 and 2 in Figure 2-C. At site 1, a deformation of the backbone characteristic of AP sites, a bulge, is identified in all the three simulations, with an extrahelicity less pronounced than the typical 30% value in B-DNA. The cluster analysis shows that the conformational behavior of this site is rather monotonous, with a percentage of occurrence of the largest cluster of 77 %, 83%, and 99 % for the three simulations, respectively - see Figure S3. The AP site is not involved in strong interactions within the double-helix. It is interacting only transiently with the facing, orphan dA20 or the adjacent dT6 and dT8. In MD1 and MD3, structural fluctuations sometimes allow transient, weak interactions between the orphan base dA20 and dT8.

AP at site 2 exhibits a much more complex structural behavior. One can visualize on Figure 2-D the two representative conformations of the main clusters identified from the simulations. Although the main cluster (64.0% of occurrence) shows the AP site in its inchworm conformation, the second one reveals a B-DNA-like structure triggered by the rotation of the initially ejected AP and dA5 within the double-helix. In MD1 and MD3, the ejected adenine can flip back within the double helix - see Figure S3. In MD1, the two most representative clusters (43 % and 35% of occurrence) indeed show a marked return of the ejected adenine to its initial position within the major groove. Similarly, dA5 starts to come back within the major groove in the last 100 ns of MD3, corresponding to the third cluster (11%). The latter could be more populated if the trajectories were spanning several microseconds, allowing sufficient time for dA5 to fully retrieve its canonical positioning in the double helix. Interestingly, the extrahelical character of the AP site seems to be positively correlated to the rotation of dA5 out of the helix - see Figure S4. The evolution of the distance between centers of mass of dA5 and its complementary nucleobase (dT22) was chosen as a descriptor of dA5 ejection from the double helix along the trajectory. The threshold distance corresponding to dA5 ejection was fixed at a lower bound of 10 Å by observation of the MD conformational ensembles, the reference distance in canonical B-DNA lying at 6.8 Å - see Figure S4. The evolution of the AP site extrahelicity and dA5 ejection along the MD trajectories indeed shows that the adenine dA5 movement back within the double helix is promoted by a decrease of AP extrahelicity. In MD1, the overall extrahelicity of the AP site rapidly decreases in the first 100 ns, which allows dA5 to efficiently flip back within the major groove and form stable canonical interactions until the end of the simulation. In MD3, we observe the same trend in the last 100 ns with AP site extrahelicity dropping and dA5 starting to rotate towards the major groove, as can be seen on the representative structure of the corresponding cluster. The so-called inchworm conformation of AP at site 2, as found in the crystal structure for THF, remains strongly pronounced in MD2 and in the first 350 ns of MD3 gathering most of the population of cluster 1 (48%).

### PCA analysis

To overcome the tediousness and potential lack of reproducibility of the visual inspection of the AP sites with the nucleosomal DNA environment, we performed PCA on the whole nucleosome core for each MD (see Figure 3). The overall picture highlights the localization of salt bridges within the histone core or between histones and DNA (see Figure S6). The motion of the histones tails is also captured. Consistently with our previous observations, the inchworm conformation around site 2 is the most important DNA region with respect to PCs, with a predominance of the ejected dA5 and its neighbours. Its relative importance is higher for MD1 and MD3 where dA5 can flip back in the double helix. On the contrary, the contribution of the surrounding region of site 1 is low in all MD (but slightly higher in MD1). However, the contribution of the abasic site itself is similar in both sites (1.9 to 3.6 %), suggesting that its flexibility is not highly correlated to its initial conformation. These observations on the DNA sites can be correlated with the behaviour of H2A and H2B N-term tails: H2A interacts with the abasic site and does not contribute a lot to PCs whereas the H2B tail, in close contact with dA5 and the inchworm region, present a relatively high mobility. However, the per-residue importance of these tails differs depending on the MD simulation and probably the interaction between its numerous positively-charged residues and the DNA.

**Figure 3.**
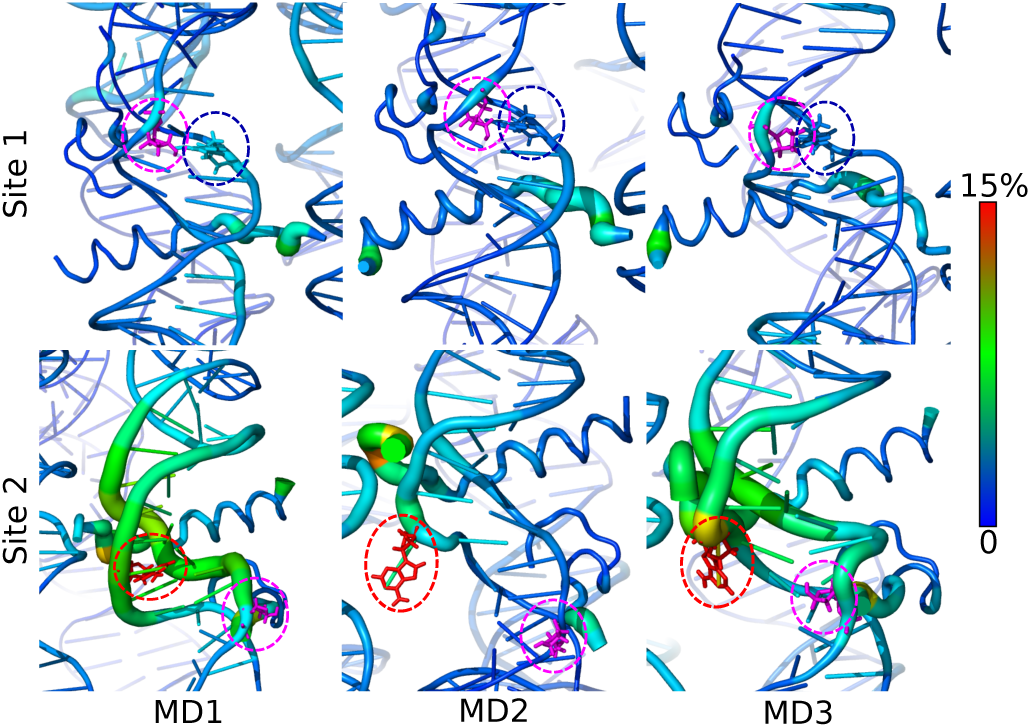
Cartoon representation of the per-residue relative contribution to the 10 first principal components around site 1 (top) and site 2 (bottom), detailed for the three MD replicates. Only DNA and the N-term tails of H2A and H2B are represented. Abasic sites, dA5 for site 2 and dA20 for site 1 are depicted in magenta, red and blue sticks, respectively, and encircled with corresponding colors. For a representation on the full NCP, see Figure S6.

**Figure 4.**
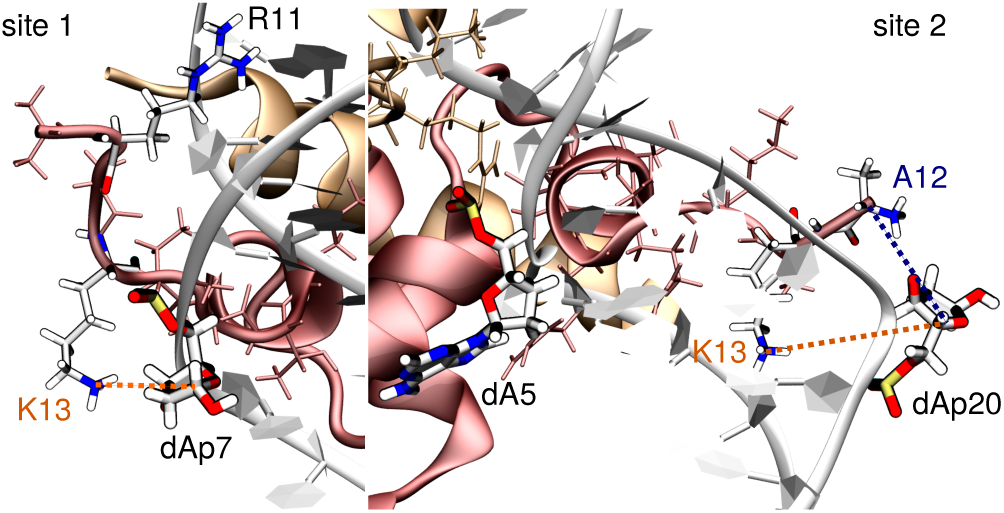
Main interactions between H2A residues and the damaged site (here AP) with K13 near site 1 (left) and K13 and N-terminal A12 near site 2 (right). H2B appears in orange and H2A in red, with their tail residues in thin licorice. The abasic sites (dAp7 at site 1 and dAp20 at site 2), the ejected adenine at site 2 (dA5), and the interacting tail residues (K13, N-term A12, and R11) are depicted in thick licorice, colored by element. We clearly distinguish R11 and K13 insertion in the minor groove two base pairs above site 1 and near site 2, respectively.

### Stabilization of the inchworm conformation for THF derivatives

Experimentally, tetrahydrofuran (THF, see Figure 1-A) is often invoked as an experimental model for deoxyribo-type AP sites, since this analog offers a higher stability (10, 15, 40). Yet it presents chemical and structural differences, with consequences in terms of repair (41). Simulations for the THF-type AP-containing nucleosome were performed to analyze the structural behavior of THF within a NCP. Unlike AP sites, THF moieties lack the chemical features to form hydrogen bonds within the double helix. As a consequence, they tend to show a much higher extrahelicity than canonical AP sites in both the NCP and the reference 15-bp oligonucleotide - see Table 1. At site 1, the THF-type AP lesion remains within the double helix, often slightly rotated towards the major groove - Figure S3. No significant reorganization of the surroundings is observed, with the facing dA20 only transiently interacting with dT6 and dT8 flanking THF. Contrary to the control 15-bp, its extrahelicity is found to be very low, i.e. 5.3% in average.

At site 2, the structural behavior is also much more stable than for the classical deoxyribo-type AP site. The inchworm conformation is highly conserved in both MD2 and MD3, with 99.5% and 99.7% of extrahelicity. dA5 remains ejected, and interacts with the H2B tail as seen with deoxyribo-AP - see Figure S5. As dA5 and THF remain ejected from the double helix, a reorganization of the surrounding nucleobases is observed. A strong Watson-Crick pairing and *π*-stacking is formed, as could be found in a standard B-DNA conformation. In MD1, however, THF tends to slightly rotate back within the double helix. As a consequence, its extrahelicity drops to 56.6%. Interestingly, the lower extrahelical character of THF does not favor dA5 rotation back within the helix as can be observed with AP sites - see Figure S3.

### Interaction with histone tails

Beyond the structural distortion of the DNA duplex, the nucleosomal environment offers a high number of residues prone to interact with the phosphate backbone, as well-known, but also and perhaps competitively with the abasic site and the orphan nucleobase. Our simulations reveal the mobility of the proximal H2A and H2B N-term tails to the damaged sites located at SHL 4.5, as highlighted by the PCA analysis - see Figure 3. In-depth inspection of histone tails interactions shows that positively-charged lysines and N-term residues are close to damaged sites, prone to react with AP site towards the formation of DNA-protein cross-links.

Notably, H2A’s K13 is found within 6 Å of both damaged sites in 6.0-9.2% and 12.2-12.8% of the total simulation time with AP and THF, respectively - see Table 2. Interaction between AP20 and the positively-charged N-term A12 amino acid of the truncated H2A tail is also observed (see Figure 4), more frequently with the canonical AP site (~20%) than with its THF analog (~7.3%). The H2A tail interacts most of the time in the minor groove nearby both sites, often through insertion of a positively-charged residue, typically R11 two base pairs above site 1 and K13 near site2.

**Table 2.** Averaged percentage over the 3 replicates of residence of the charged amino groups in the surroundings of the damaged site (AP or THF) and with an approaching distance inferior to 6 Å.

| Interaction | AP (all) | AP (MD1) | AP (MD2) | AP (MD3) |
| --- | --- | --- | --- | --- |
| K13 - AP7 | 6.0 % | 4.5 % | 6.1 % | 7.5 % |
| K13 - AP20 | 9.2 % | 8.3 % | 7.9 % | 11.3 % |
| A12 - AP20 | 19.9 % | 5.0 % | 30.0 % | 24.8 % |
| Interaction | THF (all) | THF (MD1) | THF (MD2) | THF (MD3) |
| K13 - THF7 | 12.8 % | 1.8 % | 6.2 % | 6.6 % |
| K13 - THF20 | 12.2 % | 4.8 % | 14.5 % | 17.4 % |
| A12 - THF20 | 7.3 % | 3.2 % | 10.6 % | 8.2 % |

The ejected orphan adenine at site 2 (dA5) also interacts with histone tails, but mostly with H2B and the backbone of the second turn double-helix of DNA wrapped onto the histone core - see Figure S5. In simulations with AP sites, these interactions fade as the adenine slides back within the DNA, which is not observed with THF as the inchworm conformation is conserved. This suggests that the presence of AP sites instead of THF destabilize the inchworm conformation by favoring the B-DNA like structure.

### Comparison with the control 15-bp sequences reveals a particular sequence effect

In the 15-bp oligomer, the dA-dT base pairs surrounding the abasic site tend to swing among each other. It triggers the reorganization of the internal Watson-Crick pairing as the abasic or THF site is ejected and re-inserted within the double helix along the simulations. The THF analog exhibits a much more labile behavior than the canonical AP, characterized by a much higher extrahelicity of 42.7% and 72.8% for site 1 and site 2 sequences, respectively, against the usual 30-40% for AP - see Table 1. Interestingly, the inchworm conformation is observed for site 1 reference sequence with AP, with ejection of an adenine corresponding to the one that is also excluded in front of site 2 in the crystal. As the artificial 145-bp DNA sequence from Osakabe and coworkers was chosen as an alternation of dA-dT and dC-dG tetramers, the reorganisation of the dA-dT pairing around the lesion site stays easy. As such, ejection of one of a nucleobase on the facing strand, concomitantly to an extrahelical lesion site, allows the stacking of the remaining trimer in a canonical B-DNA shape, as was observed for clustered abasic sites in previous works (15, 21).

## DISCUSSION

In direct line with the recent experimental X-ray investigation by Osakabe and coworkers, (12), our simulations allow to test at the microsecond range the stability on the so-called “inchworm” conformation of tetrahydrofuran-type abasic sites. THF-type abasic sites are confirmed to be stable along our simulations, which rules out a possible crystal packing effect.

A direct comparison between canonical vs. THF-type abasic sites is absent from the literature, especially within a NCP, yet THF analogs are often invoked experimentally to mimic AP site (10, 40). This MD investigation allows to “restore” a deoxyribo-type abasic site with exactly the same nucleosomal embedding. It also offers a same-footing comparison with control B-DNA 15-bp oligonucleotides. Our simulations show that the nucleosomal environment reshapes the conformational behavior of the AP site-bearing double helix with respect to the control naked DNA, by imposing structural constraints through strong interactions with the histone octamer. These structural constraints are reflected in the value of the bend angle around the histone core (42). The bend angle of the damaged NCP oligonucleotide lies at around 55^*◦*^, slightly superior to the characteristic value of 41.3° calculated at SHL 4.5 from an undamaged NCP crystal structure (PDB ID 1KX5). Interestingly, NCP embedding vanishes the differences of bend angle between the canonical AP and its THF analog - see Figure S2.

Formation of an abasic site leaves a “hole” in the double-stranded helix, which can evolve towards flipping out of the AP site or of the orphan nucleobase. The structural outcome of the orphan base, here an adenine, is highly driven by sequence effects (21). The sequence used by Osakabe et al. is a succession of dG-dC and dA-dT tetramers. Consequently, the ejection of the AP-site leads to the rotation of an adenine on the facing strand to allow the stabilization of the tetramer in a canonical trimer-like conformation. This reorganization is observed at site 2 in NCP, but also in the control 15-bp oligomers simulations, independently of the lesion site for which they are symmetrical. In previous work, we already observed this kind of behavior upon presence of clustered abasic sites and THF in control B-DNA (15, 21). Osakabe et al. suggested that site 1 cannot exhibit the inchworm conformation because of the stretching of the double helix around the histone core induced the reorganization around the ejected lesion at site 2. In our simulation, site 1 indeed remains in a B-DNA-like conformation, with the facing dA20 orphan. Nevertheless, while THF extrahelicity at site 2 remains high along the simulations, highlighting an inchworm-like conformation, deoxyribo-AP site showed a different structural behavior. Our simulations revealed that the AP site tends to rotate back within the double helix, concomitantly with the ejected dA5 flipping back to its canonical position. According to our PCA analysis, these motions in site 2 come with a slightly more dynamical site 1. The propensity of dA5 to flip back within the double-helix appears to be favored by extrahelicity drop of the facing lesion (see Figure S4), but also to its interaction with the histone core. Within THF-harboring NCP, the ejected adenine interacts strongly with the H2B tail and the backbone of the second turn DNA helix of the NCP (Figure S5), the inchworm conformation of THF at site 2 remaining conserved in the simulations. On the contrary, these interactions are destabilized upon presence of an AP site instead of THF. The additional hydroxyl group of the AP site might enhance interactions in its surroundings. Therefore, by promoting the classical B-DNA conformation against the inchworm one, the presence of the AP site might trigger the destabilization of dA5 initial interactions to favor its rotation back within the double helix - see Figure S5. As revealed by the PCA analysis, the contribution of the adenine in the overall motion of the system is much more pronounced for the simulation in which site 2 re-adopts a classical B-DNA conformation (MD1 and MD3), favored by the disruption of dA5 initial interactions with H2B and the closeby DNA backbone.

Interactions between histone tails and abasic sites are also a most important to seek, for they can lead to the formation of DNA-protein cross-links via a Schiff base intermediate between AP and lysines or N-terminal residues, ultimately resulting in single strand breaks. It has been established experimentally that H2A is responsible of ~ 77% of the DPCs with AP sites at SHL 4.7 (43), together with H2B ~90%. We indeed observe the close proximity of H2A K13 to the damaged site in the MD simulations. THF and AP sites are within 6Å of either K13 or the positively-charged N-term A12 during up to ~20% of the simulation time. Our simulations provide an accurate measure of the contact occurring between AP sites and positively-charged residues. This could be generalized to other nucleosomes, notably more canonical sequences such as *α*-satellite. Indeed the AP-containing nucleosomal structure obtained by X-ray (12) features a particular repetition of tetramers which was chosen to promote crystallization, but is somehow less representative of canonical N-DNA. Our simulation protocol restores a dynamical view of AP-containing DNA in interaction with a histone core, which remained elusive from the literature. We are currently using the same protocol to explore the dynamics of NCP in the presence of clustered abasic sites, to rationalize the drastic increase in DPCs formation rates compared to single AP-sites (44). MD simulations provide important insights into the finely-tuned sequence effects and dynamics of AP-sites within NCP, which are of utmost important to unravel the complicated mechanisms of their repair (45).

## ACKNOWLEDGEMENTS

The authors thank the SYSPROD project and AXELERA Pôle de Compétitivité for financial support (PSMN Data Center). V.C. is grateful for an ATOSIM Master research grant. This work was performed within the framework of the LABEX PRIMES (ANR-11-LABX-0063) of Université de Lyon, within the program ‘Investissements d’Avenir’ (ANR-11-IDEX-0007) operated by the French National Research Agency (ANR).

