## Supplementary material for "Nucleosomal embedding reshapes the dynamics of abasic sites": S

§*Current address: Université Côte d'Azur, CNRS, Institut de Chimie de Nice UMR7272, Nice 06108, France*

### Force field parameters and charges for THF

Force field parameters and charges for the 1',2'-dideoxyribofuranose-5'-phosphate nucleotide (THF) were generated using the same protocols as previously applied for AP-sites (see Bignon *et al.* NAR 2016,44, 8588–8599). The atom type assignment appears in Table S1. As all parameters were already available in the ff14SB Amber force field for these atom types, no additional parameters had to be generated.

Table S1: Numbering, names, types, and charges of THF atoms.

| Atom number | Atoms name | Atom type | Atom charge |
|---|---|---|---|
| 1 | P | P | 1.166 |
| 2 | OP1 | O2 | -0.776 |
| 3 | OP2 | O2 | -0.776 |
| 4 | O5’ | OS | -0.495 |
| 5 | C5’ | CJ | -0.007 |
| 6 | H5’ | H1 | 0.075 |
| 7 | H5” | H1 | 0.075 |
| 8 | C4’ | CT | 0.163 |
| 9 | H4’ | H1 | 0.118 |
| 10 | O4’ | OS | -0.448 |
| 11 | C1’ | CT | 0.057 |
| 12 | H1’ | H1 | 0.079 |
| 13 | H1” | H1 | 0.079 |
| 14 | C3’ | C7 | 0.071 |
| 15 | H3’ | H1 | 0.098 |
| 16 | C2’ | CT | -0.126 |
| 17 | H2’ | HC | 0.084 |
| 18 | H2” | HC | 0.084 |
| 19 | O3’ | OS | -0.523 |

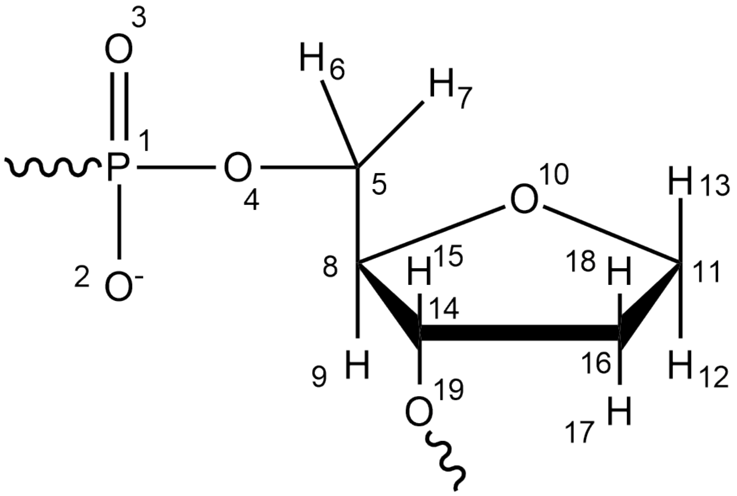


Figure S1: Chemdraw representation of the 1’,2’-dideoxyribofuranose-5’-phosphate nucleotide and corresponding atom numbers.

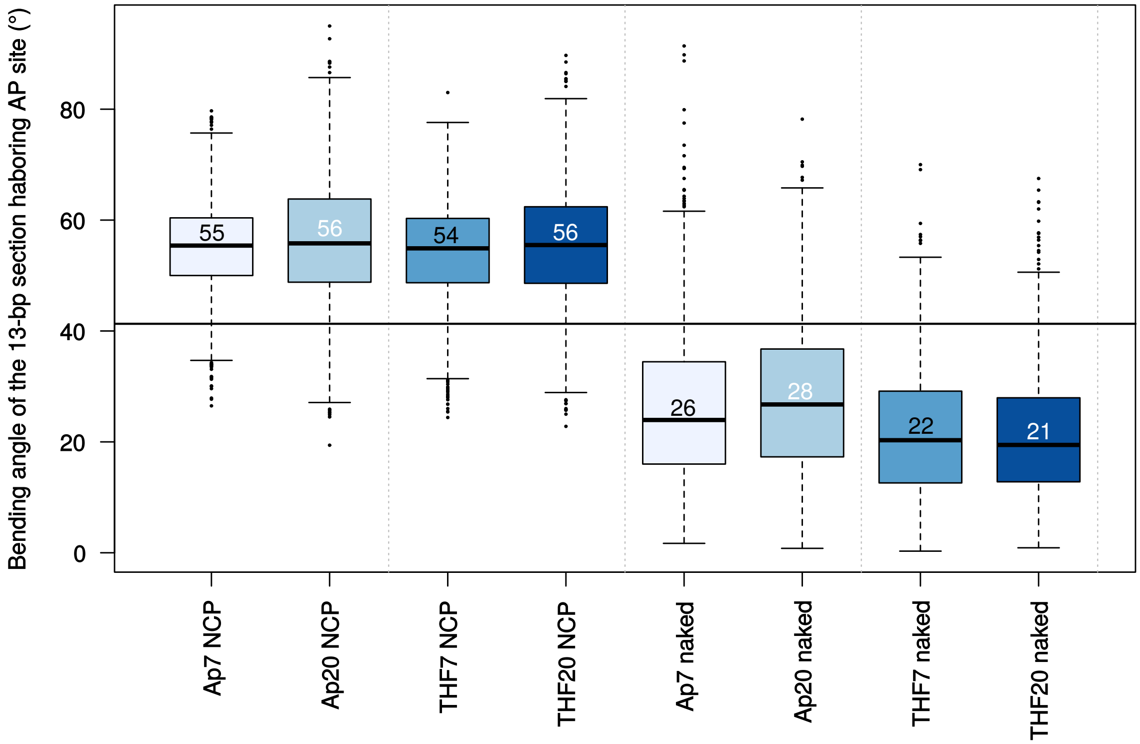


Figure S2: Bending angle of the 13-bp harboring the lesion site , for AP and THF at sites 1 (Ap/THF7) and 2 (Ap/THF20). Averaged values over the three replicates of damaged NCP appear on the left and bend angles of control DNA oligonucleotides are displayed on the right. The black horizontal line corresponds to the reference value calculated from the 1kx5 crystal structure of an undamaged NCP at SHL4.5 (41°).

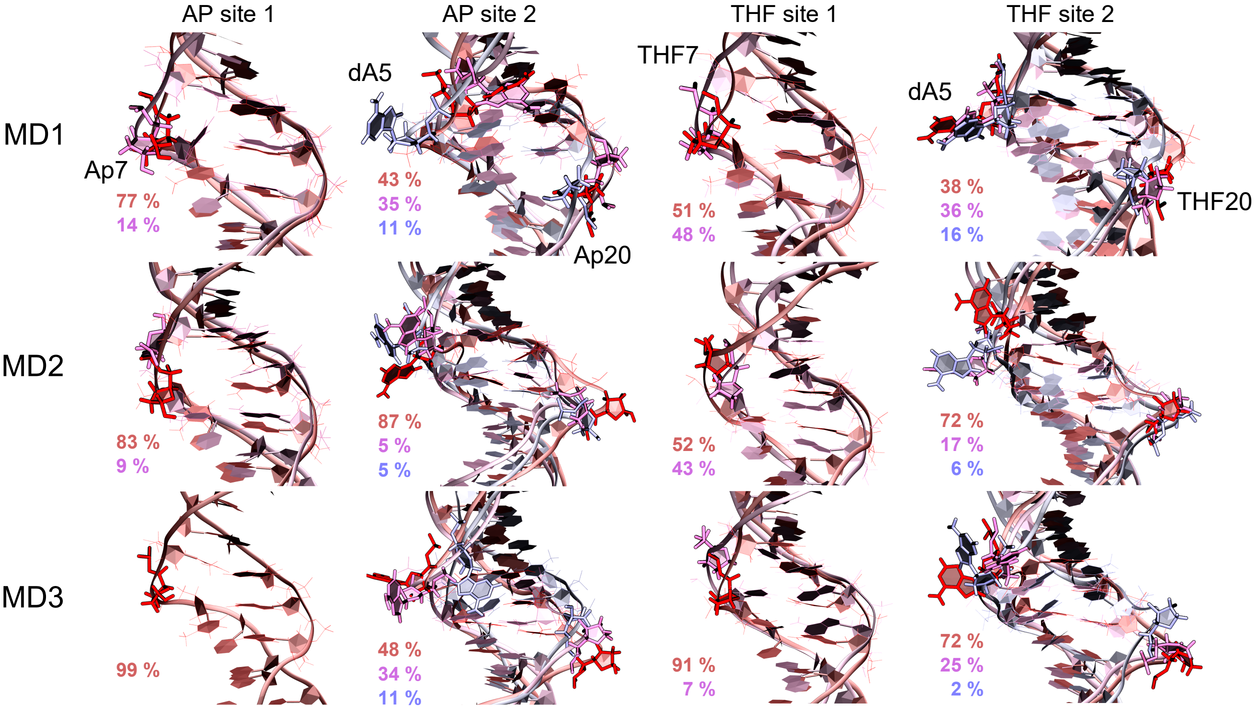


Figure S3: Representative structures of the three main clusters of damaged site 1 and site 2 for AP (left) and THF (right) within the nucleosome. Percentages of occurrence of the major clusters are detailed for each of the three replicates (MD1, MD2, MD3). The most important cluster appears in red, the second one in pink, and the third one in pale blue. The color code used for the percentages is the same. In case the second and third clusters are negligible, only the first one is showed (eg., for AP site 1 MD3).

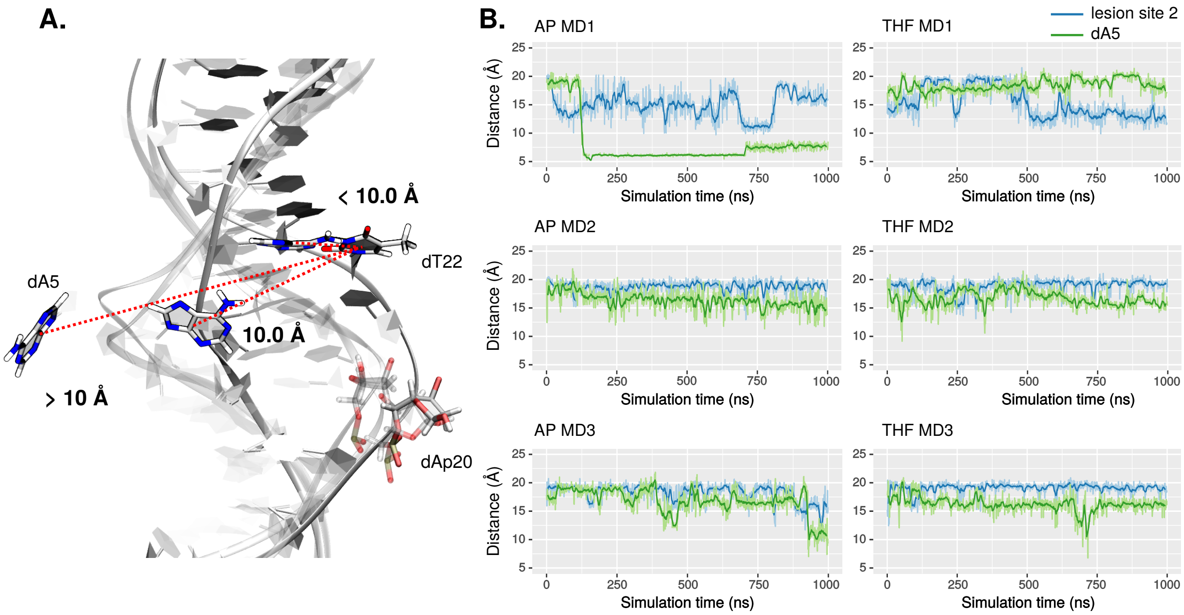


Figure S4: A. dA5 extrahelicity states based on the distance between dA5 and its paired dT22 nucleobases centers of mass. The 10 Å threshold was defined from observation of MD trajectories, the nucleobase being considered extrahelical above this value. In canonical B-DNA, the value of this distance is found around 6.8 Å. B. Extrahelicity of the lesion site 2 and dA5, for the three MD replicates with AP site (left) and with the THF analog (right)

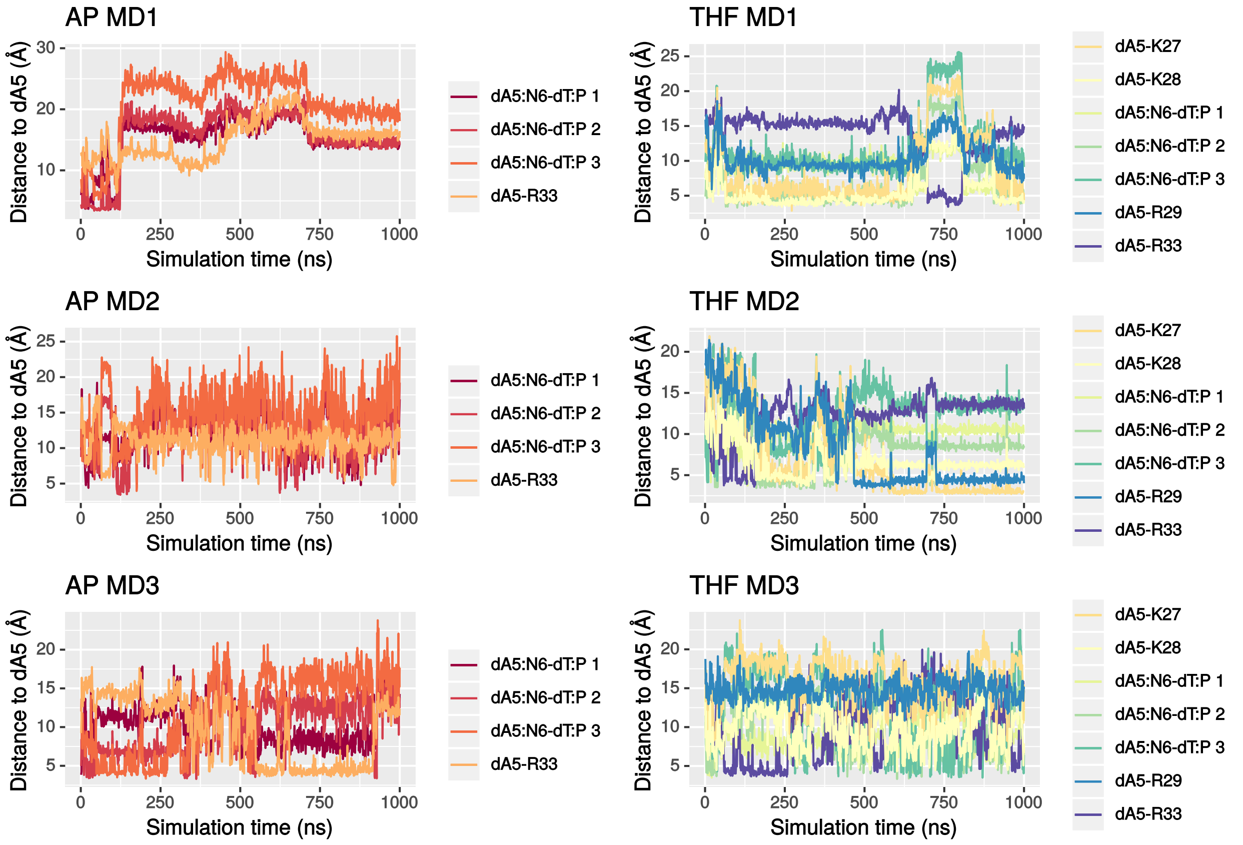


Figure S5: Distances of surrounding DNA backbone phosphates (dT:P 1, 2 and 3) and H2B residues (lysines and arginines) to the ejected dA5 in the three replicates with AP (left) and THF (right).

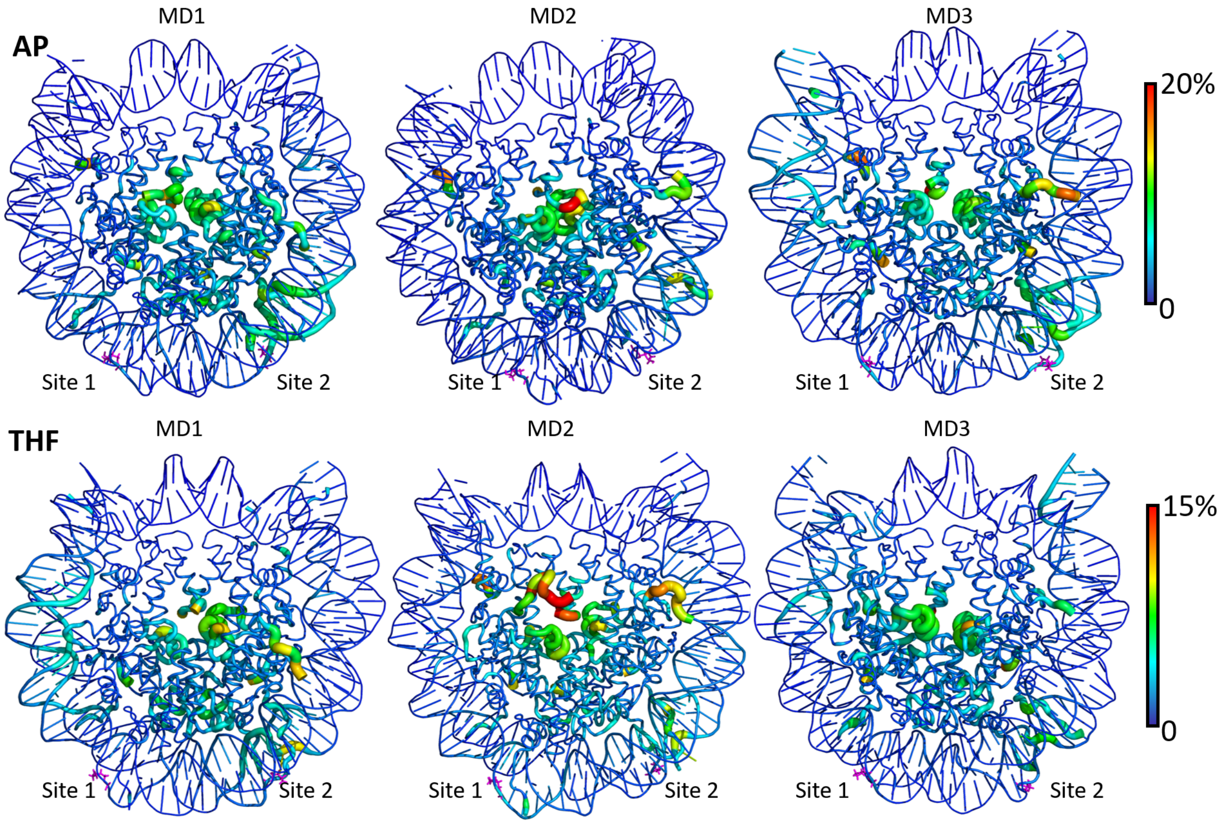


Figure S6: Cartoon representation of the per-residue relative contribution to the 10 first principal components for each replicates of AP- and THF-containing NCP systems. Abasic sites are depicted in magenta sticks.
